# Mutations generated by repair of Cas9-induced double strand breaks are predictable from surrounding sequence

**DOI:** 10.1101/400341

**Authors:** Felicity Allen, Luca Crepaldi, Clara Alsinet, Alexander J. Strong, Vitalii Kleshchevnikov, Pietro De Angeli, Petra Palenikova, Michael Kosicki, Andrew R. Bassett, Heather Harding, Yaron Galanty, Francisco Muñoz-Martínez, Emmanouil Metzakopian, Stephen P. Jackson, Leopold Parts

**Affiliations:** Wellcome Sanger Institute, Wellcome Genome Campus, Hinxton, Cambridgeshire, CB10 1SA, United Kingdom; Cambridge Institute of Medical Research, University of Cambridge, Wellcome Trust MRC Building lab 6.36, Hills Road, Cambridge, CB2 0XY, United Kingdom; The Wellcome Trust/Cancer Research UK Gurdon Institute and Department of Biochemistry, University of Cambridge, Cambridge CB2 1QN, United Kingdom; Department of Computer Science, University of Tartu, J. Liivi 2, 51001, Tartu, Estonia

**Author notes:** Correspondence to LP or FA. These authors contributed equally to this work (Felicity Allen, Luca Crepaldi).

## Abstract

The exact DNA mutation produced by cellular repair of a CRISPR/Cas9-generated double strand break determines its phenotypic effect. It is known that the mutational outcomes are not random, and depend on DNA sequence at the targeted location. Here, we present a systematic study of this link. We created a high throughput assay to directly measure the edits generated by over 40,000 guide RNAs, and applied it in a range of genetic backgrounds and for alternative CRISPR/Cas9 reagents. In total, we gathered data for over 1,000,000,000 mutational outcomes in synthetic constructs, which mirror those at endogenous loci. The majority of reproducible mutations are insertions of a single base, short deletions, or long microhomology-mediated deletions. gRNAs have a cell-line dependent preference for particular outcomes, especially favouring single base insertions and microhomology-mediated deletions. We uncover sequence determinants of the produced mutations at individual loci, and use these to derive a predictor of Cas9 editing outcomes with accuracy close to the theoretical maximum. This improved understanding of sequence repair allows better design of editing experiments, and may lead to future therapeutic applications.

## Introduction

CRISPR/Cas9 is a transformative DNA editing technology (Doudna and Charpentier 2014). It operates by recruiting the Cas9 nuclease to a genomic locus with a protospacer-adjacent motif (PAM) using a short synthetic guide RNA (gRNA) with a 18-20 nt sequence matching the desired target. Cas9 then cuts DNA at that location, and when the double strand break is repaired by cellular machinery, frameshift mutations can occur, disabling translation of the correct protein. This enables genome-wide CRISPR/Cas9 screens in mammalian cells, in which every protein coding gene is individually knocked out to assess its impact (Doench 2018; Wang et al. 2014; Shalem et al. 2014; Koike-Yusa et al. 2013). While such experiments allow high-throughput interrogation of nearly all facets of cellular function, fundamental aspects of the editing process remain poorly understood, central of which is the profile of produced mutations.

Cas9-generated mutations result from imperfect action of DNA repair pathways that are activated to remedy the double strand break. The major repair mechanisms include non-homologous end joining, which re-ligates the generated ends, often introducing errors of a few nucleotides; and microhomology-mediated end joining, in which short tracts of local matching sequence anneal, ultimately resulting in deletion of the intervening bases (Chiruvella, Liang, and Wilson 2013; Her and Bunting 2018). Less frequently, the homology-directed repair pathway uses intact sequence from the homologous chromosome to more faithfully fill the gap at the break. Choice of pathway is influenced by a host of factors including cell cycle stage, and availability of repair enzymes (Truong et al. 2013). Perhaps surprisingly then, the frequency of alternative Cas9 editing outcomes (“the mutational profile”; Figure 1) is largely reproducible, and depends on the targeted sequence (Bae et al. 2014; van Overbeek et al. 2016; Koike-Yusa et al. 2013; Lemos et al. 2018), indicating that the errors in repair occur in a non-random manner. While DNA repair pathways and their key components have been characterized, the biases that favour one mutation over another are not fully understood, especially for the breaks inflicted by Cas9.

**Figure 1.**
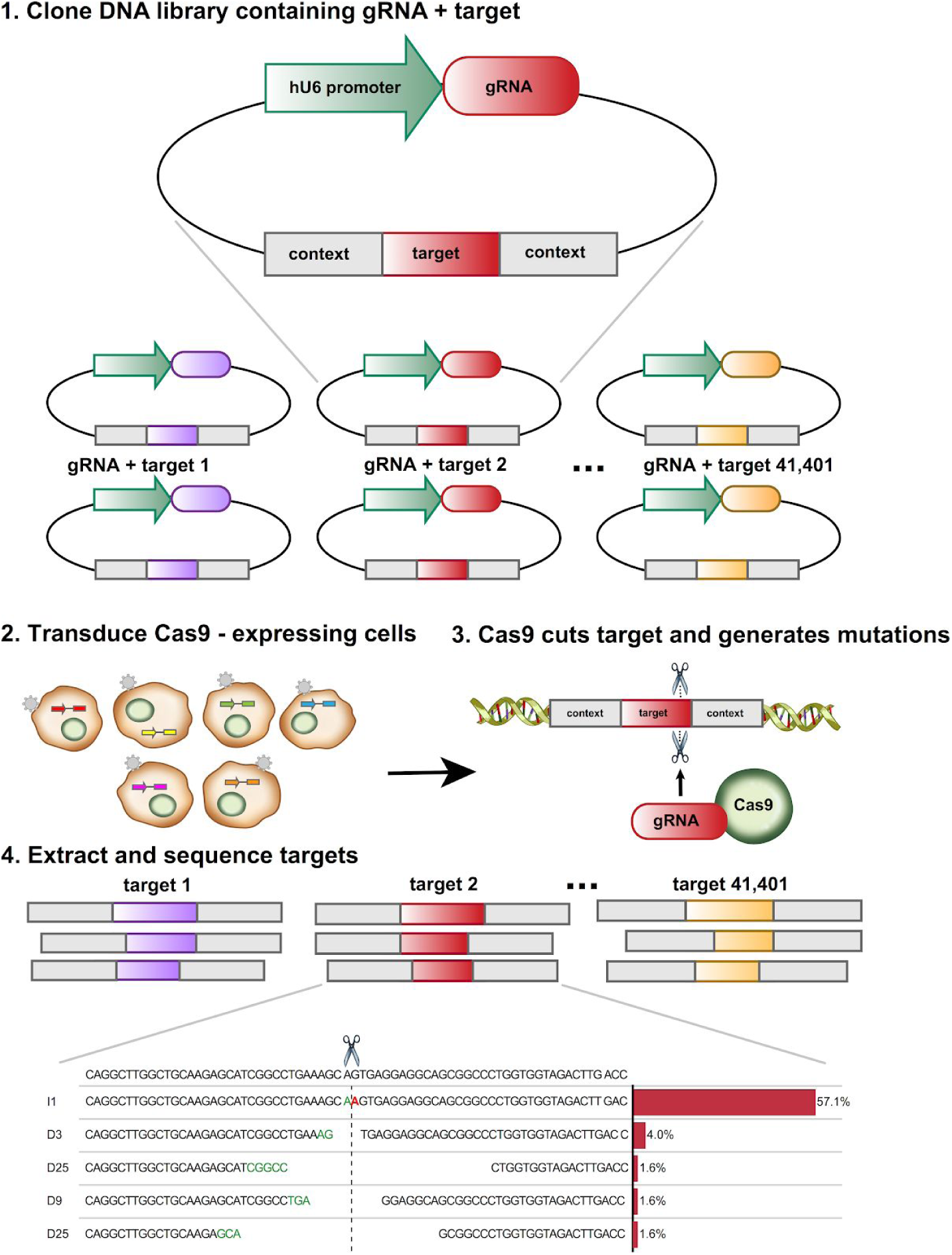
Mutational profiles generated by CRISPR/Cas9, and a method for their high-throughput measurement. High throughput measurement of repair outcomes. Constructs containing both a gRNA and its target sequence (matched colors) in variable context (grey boxes) are cloned en masse into target vectors containing a human U6 promoter (green) (1), packaged into lentiviral particles, and used to infect cells (2), where they generate mutations at the target (3). DNA from the cells is extracted, the target sequence in its context amplified with common primers, and the repair outcomes determined by deep short read sequencing (4).

Mutations generated by Cas9 determine the effect of editing on the targeted gene. Despite their importance, they are not directly observed during CRISPR/Cas9 screening, and are yet to be studied systematically. No robust predictive model for the mutations created by each gRNA exists, and current methods for screen design (Doench et al. 2016) and analysis (Li et al. 2014; Allen et al. 2018; Hart and Moffat 2016) do not fully account for the diversity of functional outcomes. This lack of information also precludes use of a non-templated Cas9 editing approach for therapeutic purposes, for which a particular outcome may be preferred, but whether a gRNA can produce it cannot be predicted.

To date, mutational profiles have not been measured at scale. The main barrier has been the labour necessary to individually amplify the sequence at each of the targeted loci. The largest dataset of genomic repair profiles to date comprises 436 profiles examining 96 unique gRNA sequences using the Cas9 protein from *S. pyogenes* (van Overbeek et al. 2016). More gRNAs (∼1400) were employed in a study that introduced the target and gRNA into cells simultaneously (Chari et al. 2015), but the low probability of a gRNA and its corresponding target meeting in the same cell resulted in an average mutation rate of 0.2%, and insufficient data for a comprehensive analysis. A synthetic approach introducing target in the same construct was used for the Cpf1 nuclease (Kim et al. 2016), which has a shorter scaffold sequence, enabling a simpler library cloning procedure to assess more gRNAs. However, the properties of the DNA breaks generated by Cpf1, as well as of the protein itself, mean that these result are not applicable to Cas9.

Here, we present the first large-scale measurement of Cas9-generated gRNA repair profiles. We synthesized over 40,000 DNA constructs, each containing both a gRNA and its target, introduced them into Cas9-expressing cell lines, and sequenced the targeted loci. We confirm that our measurements are informative of events at endogenous sites, describe the dominant outcomes and their sequence dependence in a range of cell lines, and present an accurate predictive model for forecasting the outcomes of an edit.

## Results

### Measuring repair outcomes *en masse*

The main hurdle in measuring a large number of repair outcomes is the need to selectively amplify each targeted locus. To circumvent this, we designed a construct that encodes a gRNA expression cassette together with its 23nt PAM-endowed target sequence within a larger 79nt variable context, and flanked by common PCR priming sites on both sides (Figure 1, S1). The variable context allowed us to systematically change the local sequence to directly test its influence on the repair outcome, and to unambiguously assign each sequenced target to its gRNA-target pair of origin. Using high throughput oligonucleotide synthesis followed by custom cloning reactions (Methods), we generated a library of 41,362 gRNA-target pairs. We delivered these constructs into cells using lentivirus, cultured for up to 20 days to ensure completed editing, then isolated genomic DNA, amplified the target sequence in its context, and measured the frequency of insertions and deletions that occurred by sequencing. We validated our approach in human K562 chronic myelogenous leukemia (K562) cells, collecting a median of 8,958 mutated reads (after filtering and quality control - Methods) per construct across replicate experiments. We then applied the same assay in a range of diverse mammalian cell lines, and with several Cas9 varieties, using an average coverage of 500-1600 cells per construct in each experiment (Table S2, Figure S2). The data set collected provides a deep, high resolution picture of Cas9-induced double strand break repair outcomes.

### Synthetic repair profiles are reproducible, and faithfully capture endogenous repair outcomes

First, we demonstrate that our measurements are sequence-specific and reproducible in the K562 cell line. Here and elsewhere, we use the symmetric Kullback-Leibler divergence (“KL”), a natural information-theoretic metric related to relative entropy of probability distributions, to quantify similarity of outcome frequencies (Methods). Given adequate read coverage, biological replicates measuring the same gRNA target were similar (median KL=0.7; Figure 2A,C), while targets of different gRNAs had markedly different repair outcomes (median KL across randomly selected pairs=4.8; Figure 2C). This shows that the mutational profiles are reproducible, and sequence-specific. Further, the fraction of frameshift edits, a factor that is arguably most important for knock-out experiments, is also highly correlated between biological replicates (Pearson’s R=0.9; Figure 2D).

**Figure 2.**
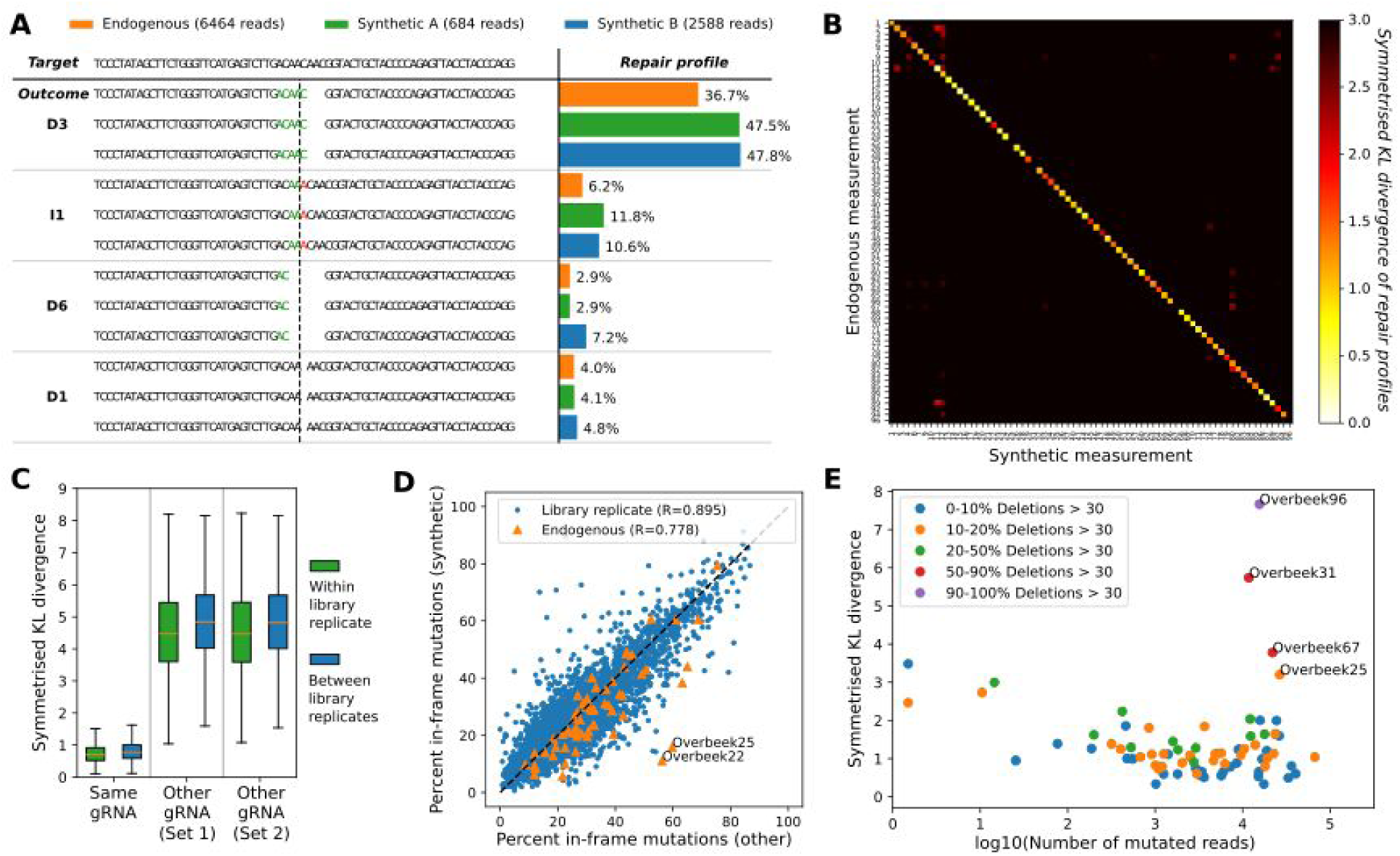
Synthetic mutational profiles are replicable, specific to individual gRNAs and closely resemble endogenously measured profiles in human K562 cells. A.Example measured repair profile reproducibility of one gRNA-target pair. DNA sequence of the target (top) is edited to produce a range of outcomes in two synthetic replicates (green, blue bars) and one endogenous measurement (orange bars). The proportions (x-axis) of the four most frequent mutational outcomes (e.g. “D3” - deletion of three base pairs, “I1” - insertion of one base pair, etc.; y-axis) is consistent between the experiments. Stretches of microhomology (green) and inserted sequences (red) are highlighted at the cut site (dashed vertical line). *B.Synthetic measurements faithfully capture endogenous outcomes. Symmetrized Kullback-Leibler divergence (white to black color scale) between the measured repair profiles (e.g. Fig. 2A) in our synthetic measurements in K562 cells (x-axis) when compared to the same individual gRNAs in endogenous measurements from van Overbeek et al. (y-axis).* *C.Synthetic measurements are reproducible and gRNA-specific. Box plots (orange median line, quartiles for box edges, 95% whiskers) of symmetrised KL divergences between replicate measurements of over 6000 gRNAs (left), as well as two different sets of randomly selected gRNA pairs (middle, right) using the same library of constructs (green boxes) and a separately cloned library of corresponding constructs (blue boxes).* *D.Frame information is reproducible between replicates, and well correlated with endogenous outcomes. In-frame percentages for one biological replicate of our synthetic measurements (y-axis) contrasted against another replicate (x-axis, blue markers, Pearson’s R=0.895), or endogenous measurements (x-axis, orange markers, Pearson’s R=0.78).* *E.Low coverage and large deletions are the main sources of discrepancy between endogenous and synthetic measurements. Symmetrized KL divergence (y-axis) between endogenous and synthetic measurements of the same gRNA editing outcomes (individual markers) is dependent on the sequencing coverage (log*_*10*_*(number of obtained reads), x-axis), and frequency of very large deletions (colors). Three target sequences that frequently give rise to very large deletions (red, purple) are not well captured by our assay by design.*

We next tested whether the measurements from our synthetic targets are a good proxy for repair outcomes at endogenous loci. To achieve this, we took advantage of data from the largest scale study of editing outcomes to date (van Overbeek et al. 2016), in which 223 human genomic targets for 96 unique gRNAs were individually amplified and sequenced (“endogenous outcomes”). Those 86 of these gRNAs that were compatible with our cloning process were included in our library with their genomic contexts. Concordance between synthetic and endogenous outcomes was very good for individual cases (Figure 2A), on the whole (median KL=1.1; Figure 2B), and for recapitulating frameshift edit fraction (Pearson’s R=0.78, Figure 2D). Nevertheless, the observed differences were larger than for biological replicates of our assay, so we next inspected the reasons for this. We identified two causes. First, sampling noise due to low sequencing coverage leads to increased divergence (Figure 2E). Second, deletions and rearrangements larger than the 30nt we can accurately measure (Methods), and which can be prevalent (Kosicki, Tomberg, and Bradley 2018), explain three of the four cases with sufficient read counts that markedly differ (KL > 3). The remaining case (“Overbeek 25”) diverges despite high reproducibility between synthetic measurements (Figure S3). Given that our construct only contains 79nt of local context, yet produces very similar outcomes for 94% of measured cases with sufficient reads (67/71), this result confirms that sequence surrounding the cut site is the major determinant of Cas9-induced mutational outcomes.

### Deletions and single base insertions are the most frequent repair outcomes in K562 cells

After concluding that our assay faithfully and reproducibly captures a majority of endogenous mutational outcomes, we further analysed a larger set of gRNA-target pairs. We selected 6,568 gRNAs from an independently designed human genome-wide library (Tzelepis et al. 2016), and measured repair outcomes in three replicates for each. Overall, we found that deletions and single nucleotide insertions were most common, with larger insertions occurring only rarely, and shorter deletions favoured over longer ones (Figure 3A). However, we also observed a long tail of larger deletion events.

**Figure 3.**
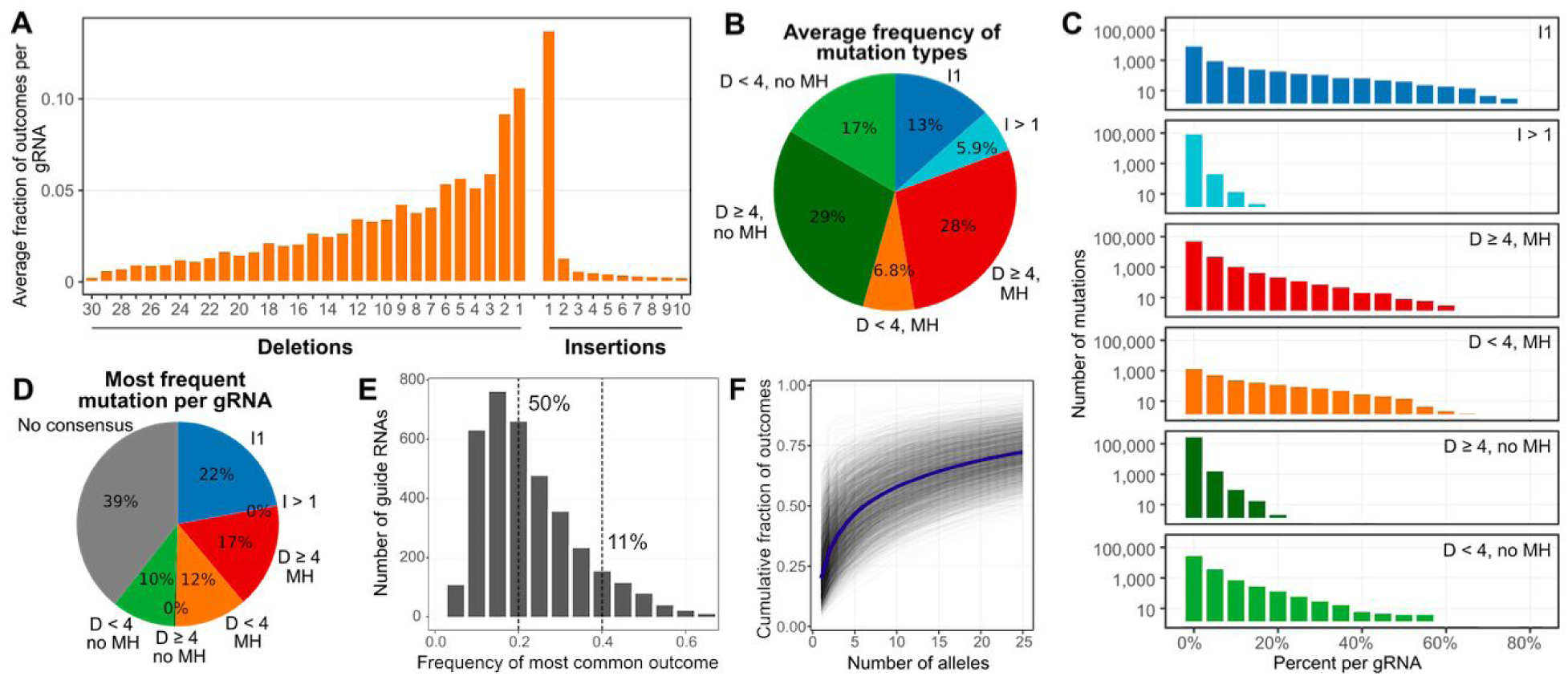
Mutational profiles are diverse and biased in K562 cells. A.Single base insertions are most common, with a long tail of moderately long deletions. The frequency (y-axis) of deletion or insertion size (x-axis), averaged across genomic sequence targets. *B.Editing outcome types are diverse. The average percent occurrence (area of wedge) of small (<4nt) and large (≥4nt) deletions, both microhomology-mediated and not, as well as small (1nt) and large (>1nt) insertions, measured in K562 cells, and averaged across genomic sequence targets.* *C.Per-gRNA event frequencies differ across indel classes. Number of individual indels* *(y-axis) as a percentage of all mutations observed for their gRNA (x-axis) separated by mutations class (rows). Colors as in (B).* *D.Specific single insertions and microhomology-mediated deletions are the most frequent reproducible mutation classes. The percent of gRNAs (area of wedge) that have the same specific indel as their most frequent mutation in all three replicates, stratified by indel class (colors). ‘No consensus’: inconsistent most frequent mutation across replicates.* *E.A single allele often accounts for a large fraction of editing outcomes for a gRNA. Number of gRNAs (y-axis) with the frequency of its most common outcome (x-axis) in K562 cells.* *F.A small number of outcomes explains most of the observed data, but many low frequency alleles are present. Cumulative fraction of observed data (y-axis) matching an increasing number of outcomes (x-axis) for each target in K562 cells (grey lines), and their average (blue line).*

We classified the repair outcomes by the type, size, and likely causal mechanism of each event. Despite shorter deletions being favoured over longer ones, most of the Cas9-generated double strand breaks (57%) resulted in a deletion of at least four base pairs (Figure 3B; cutoff defined in van Overbeek *et al.* (2016)). About half of these (28% of total) were most likely generated by microhomology-mediated end joining of at least 2nt of repeating sequence near the cut site (Methods), and the rest were likely produced by the canonical non-homologous end joining mechanism. Smaller deletions of up to three base pairs made up 23% of the observations, but are more difficult to confidently assign to a particular causal repair process. While insertion of a single base was the most common single outcome overall (13%), larger insertions were rare (6% of all mutations).

Given a similar basal activity of the different DNA repair pathways in all cells of the assay, it is natural to hypothesize that repair outcomes of individual gRNA targets largely conform to the average trend observed above. In fact, there is substantial variability in the relative frequency of different outcome types (Figure 3C). Insertions, microhomology-mediated deletions, and other deletions can all be present at frequencies ranging from near 0 to over 50% depending on the target, further highlighting the sequence-specific nature of the repair process.

Repair outcomes are biased towards particular alleles. The same specific mutation was most frequent in all three biological replicates for over 60% of gRNAs (Figure 3D), but not all mutation classes were favoured equally. The consensus mutation for each individual gRNA was almost always a single nucleotide insertion (36%), large microhomology-mediated deletion (28%), or a small deletion (36%), and could make up over half of all mutations for that gRNA (Figure 3C). In contrast, while deletions of at least 4nt without microhomology are common as a class (29%, Figure 3B), each particular event is rare, and never contributes more than 20% of the mutations (Figure 3C,D).

Overall, half of measured gRNAs have a single outcome that contributed at least 20% of the observations, and 11% have an outcome that contributed at least 40% (Figure 3E). Yet while on average, the six most frequent alleles per construct account for the majority of its observed mutations, 25 alleles collectively explain only 72% of the data (Figure 3F), indicating a large number of low frequency events. In spite of such diversity of outcomes, and the variability across gRNAs, the fraction of observations for a gRNA falling within each mutation class is highly reproducible, provided sufficient sequencing coverage (between-replicate Pearson’s R > 0.87; Figure S4). Together with evidence of profile reproducibility above, this paints a picture of a complex, yet not completely random repair process for Cas9-generated breaks.

### Repair outcomes depend on local sequence properties

The extent of microhomology between sequences on different sides of the Cas9-generated cut is known to determine some of the bias in repair outcomes (Bae et al. 2014). A large number (28,193) of our gRNA-target pairs were designed to systematically harbour a range of microhomology spans (3-15nt), separating distances (0-20nt), and mismatches (0-2nt), which enabled us to comprehensively assess their influence. We observed that the fraction of observed mutations that could be attributed to microhomology-mediated end joining was higher if the matching sequences were separated by short distances (Figure 4A). This trend held for all spans of microhomology, but was more pronounced for longer tracts (Pearson’s R=-0.7 for 10 matching bases *vs.* R=-0.2 for three matching bases; Figure 4B).

**Figure 4.**
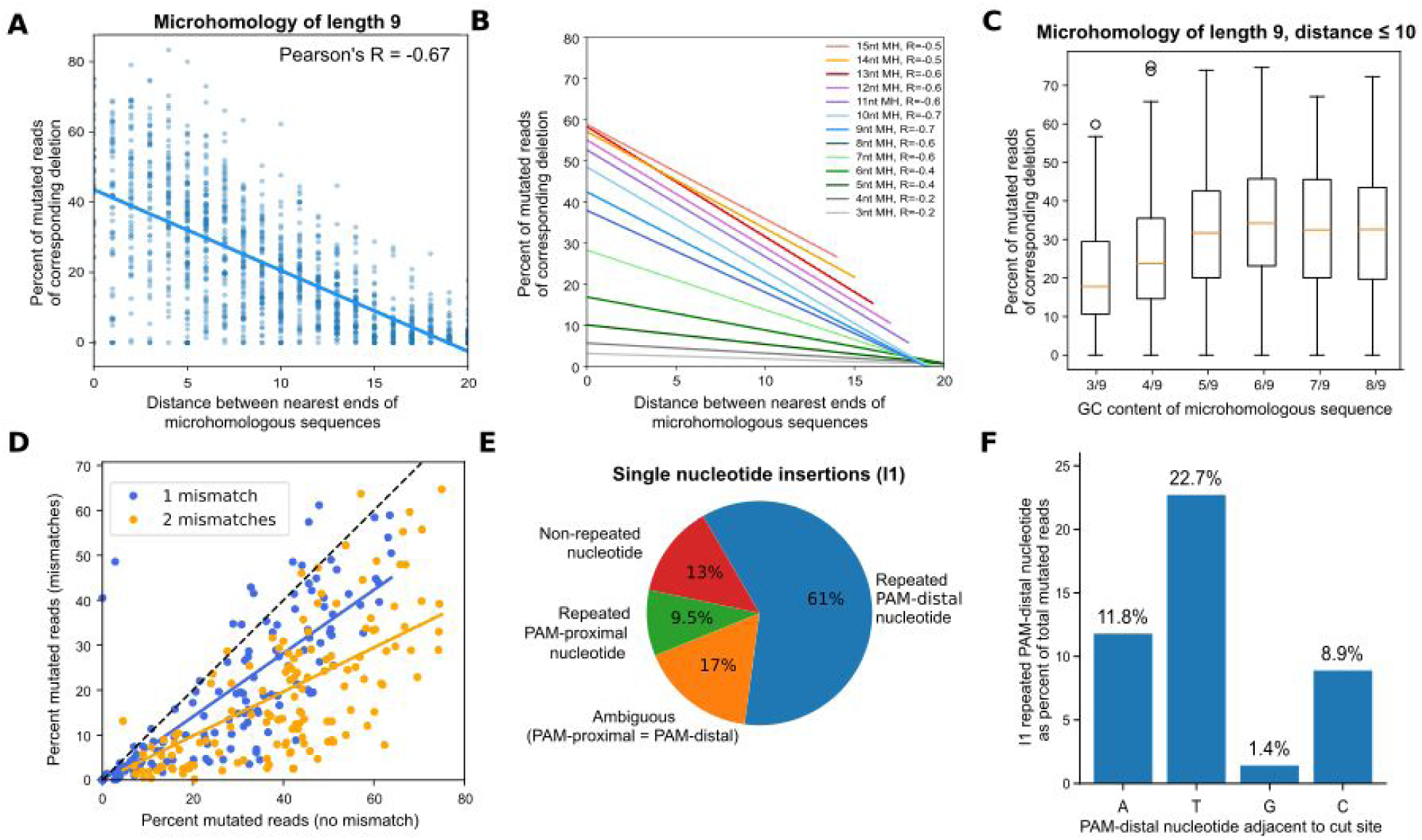
Local sequence context strongly influences editing outcomes. A.Nearby matching sequences are used as substrate for microhomology-mediated repair more frequently than distant ones. Fraction of mutated reads (y-axis) for increasing distance between matching sequences of length 9 (x-axis) for individual targets (blue markers) in K562 cells, and a linear regression fit to the trend (solid line). *B.Frequency of microhomology-mediated repair depends on the length of and distance between the matching sequences. Same as (A), but linear regression fits only for microhomologies of lengths 3 (gray, bottom) to 15 (yellow, top), with Pearson’s correlation noted in the legend.* *C.GC content influences microhomology-mediated repair fidelity. Percent gRNA reads with length 9 microhomology-mediated deletion (y-axis; boxes median and quartiles, whiskers 5% and 95%) across a range of GC contents (x-axis).* *D.Mutations in microhomology sequence reduce repair outcome frequency, but corresponding deletions are still present. For matched pairs of gRNAs, with and without mutations in the microhomologous sequence, the fraction of mutated reads associated with the particular microhomology with mismatches (y-axis) vs without mismatches (x-axis) (markers; blue: one mismatch, yellow: two mismatches; solid lines: linear regression fits). Dashed black line: y=x.* *E.The sequence context at the cut site influences which single base is inserted. PAM-distal (blue), PAM-proximal (green), and other (red) nucleotide insertion frequency of all single base insertions; if proximal and distal bases are identical (orange), no call can be made.* *F.Single base insertion rate depends on the PAM-distal base adjacent to the cut site. The average percent of per-gRNA mutations that are unambiguous PAM-distal 1bp insertions (y-axis), stratified by the PAM-distal nucleotide (x-axis).*

Sequence properties further alter the efficacy of microhomology-mediated end joining. We observed a bias due to GC content, where joining of sequences with low GC fraction (3 of 9 bases) was not observed as often as those with higher GC (Figure 4C). This could be partially explained by the sequence annealing step disfavouring the weaker A-T base pairing, analogously to preferring longer tracts of microhomology that produce a higher melting temperature for the resulting duplex. Imperfectly matching sequences were also able to generate deletions that could be attributed to microhomology-mediated end joining, but at a reduced efficiency. Indeed, the same alleles are generated at a 30% reduced rate if one mismatch is present, and 52% reduced rate if two (Figure 4D). The presence of mutations in one or the other side of the cut also allowed us to test whether one side of the sequence is preferentially retained, but we found no bias either for microhomology-mediated deletions (Figure S5) or the rest (Figure S6).

It is plausible that sequence around the Cas9-generated double strand break also influences the outcome of insertion events. Indeed, single base insertions are known to favour a repeat of the PAM-distal nucleotide adjacent to the cut site in yeast, and to an unknown extent, in humans (Lemos et al. 2018). Contrary to the lack of direction bias in deletions, we observed that in over 78% of single insertions, the PAM-distal nucleotide is inserted. Conversely, the PAM-proximal nucleotide was inserted unambiguously just 9.5%, and another nucleotide 13% of the time (Figure 4E). We further tested whether the identity of the PAM-distal nucleotide affected the rate of insertion, and found that if thymine is exposed, the double strand break is frequently repaired with insertion of another thymine, while if a guanine is present, insertions rarely occur at all (23% vs 1.4%; Figure 4F). The remaining bases are inserted at intermediate frequency (adenine: 12%, cytosine: 9%).

### Mutational outcomes vary with cell type and some Cas9 modifications

Cells can differ in activity of repair processes and/or DNA sequence, both of which influence double strand break repair outcomes (Gallagher and Haber 2018). To test whether these factors have an effect in the context of our analyses, we next performed our assay in human induced pluripotent stem cells (iPSCs), mouse embryonic stem cells (mESCs), Chinese hamster ovary (CHO) epithelial cells, as well as human retina epithelial immortalized cells (RPE-1) and leukemic near-haploid (HAP1) cell lines.

The overall distribution of repair outcome types was not the same across the different cell lines and organisms studied (Figure 5A), but the per gRNA profiles remained similar to each other (median KL < 2; Figure 5B, example in Figure S7). Some relative changes of preferred mutation classes were notable. Large insertions occurred more frequently in stem cells, with 3x and 2.1x increase in human iPSCs and mESCs, respectively, over K562 levels. The fraction of a gRNA’s mutations due to a specific large insertion generally remained below 10%, and was not reproducible in the stem cells (Figure S8), which led to increased between-replicate divergence (Figure 5B), and suggests a predominantly stochastic, sequence-independent source.

**Figure 5.**
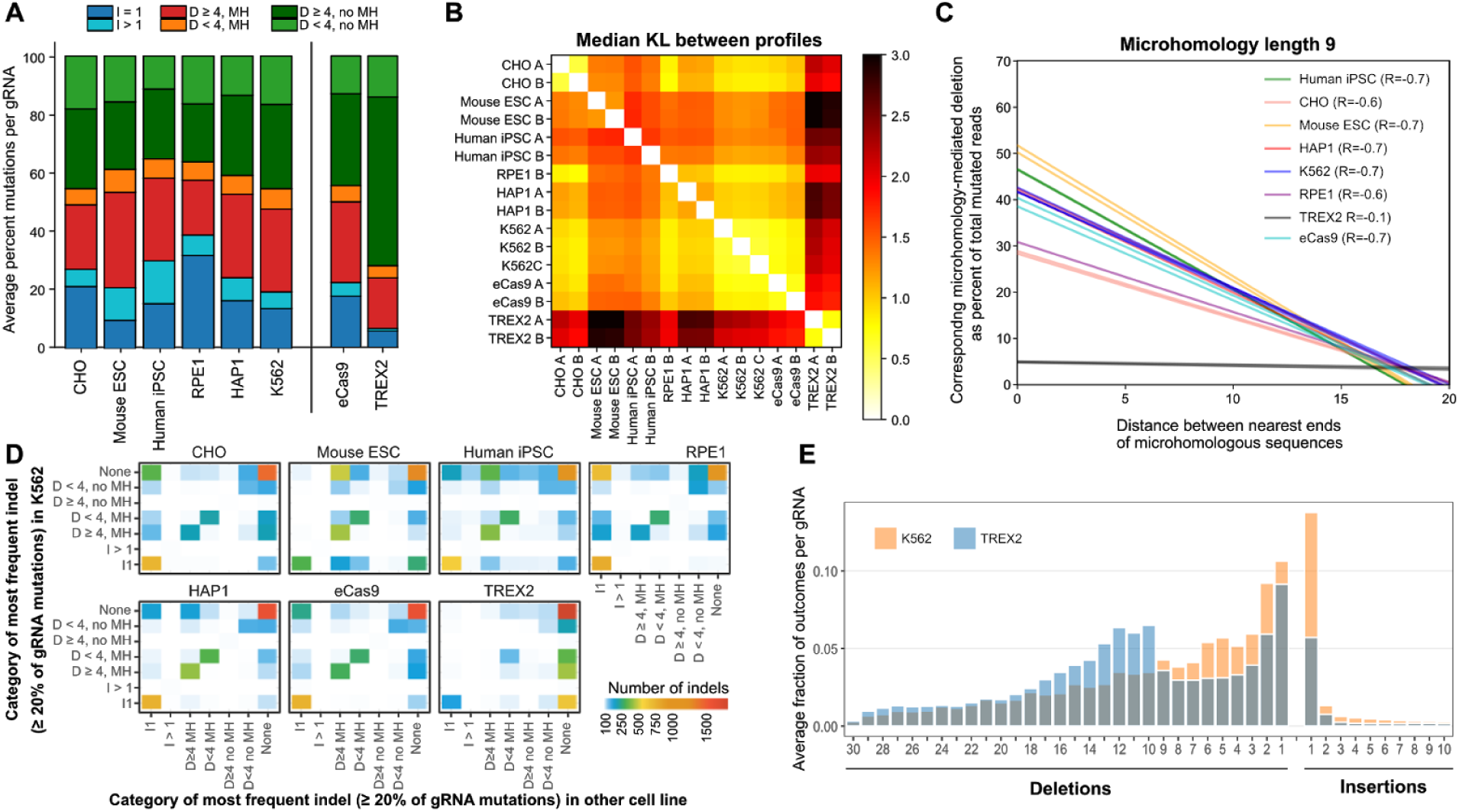
Differences between editing outcomes in K562-Cas9 and other cell lines and effector proteins. A.Genetic background influences editing outcomes. Frequency of different types of editing outcomes (y-axis; colors as 3B) for Chinese hamster ovary cell line (CHO), mouse embryonic stem cells (Mouse ESC), human induced pluripotent stem cells (iPSCs), human retinal pigmented epithelial cells (RPE-1), human near-haploid cell line (HAP1), K562 cell line, and K562 cells with alternative Cas9 proteins: enhanced Cas9 (eCas9), and Cas9-TREX2 fusion (TREX2). *B.Mutational outcomes are similar across cell lines, with consistent moderate differences in stem cells and the K562 Cas9-TREX2 fusion line. Median symmetric Kullback Leibler divergence between repair profiles (black to white color range, as in Figure 2B) in different tested lines (x and y axis).* *C.Microhomology-mediated repair fidelity is similar across genetic backgrounds, but differs for Cas9-TREX2 fusion. Regression lines (as in Fig 4A) for fraction of mutated reads* *(y-axis) for increasing distance between matching sequences of length 9 (x-axis) in K562 cells (blue) and other tested lines (colors) in multiple replicates, with overall Pearson’s correlation denoted in the legend.* *D.The type of most frequent outcome per gRNA (which is also present in above 20% for the gRNA) is mostly consistent across cell lines, but is biased towards microhomology-mediated deletions in stem cells, and I1 insertions in RPE-1 and CHO. The number of gRNAs (color) for which the most frequent indel comes from each class (x-axis) in the other cell lines examined (panels) compared to that for the same gRNA in K562 (y-axis). “None” refers to gRNAs without any indel forming more than 20% of mutations.* *E.Cas9-TREX2 fusion protein favours larger deletions compared to K562. Deletions of increasing size (x-axis) become more frequent (y-axis) in K562 Cas9-TREX2 cells (blue) compared to standard K562 Cas9 (orange).*

Deletions attributed to microhomology-mediated end joining were 38% less frequent in RPE-1 samples compared to K562; instead these cells displayed more than double the rate of single base insertion (37% in RPE-1 vs 15% in K562). The same bias was present in CHO cells, while both iPSC and mouse stem cells favoured microhomology-mediated deletions at the expense of single base insertions (Figure 5A). This trend was recapitulated in the mutation class of the most frequent indels for each gRNA, which changed depending on the cell line (Figure 5D). We found little influence of the genetic background on the link between microhomology and repair outcomes (Figure 5C), replicating our findings in K562 in other lines and species.

Multiple Cas9 effector proteins with augmented properties have been engineered, which could also give rise to changes in observed mutations. We thus considered alternative CRISPR/Cas9 reagents in K562 cells, both enhanced Cas9 eSpCas9(1.1) (eCas9) (Slaymaker et al. 2016)) and Cas9 fused to Three Prime Repair Exonuclease 2 (TREX2), which is known to increase deletion size (Chari et al. 2015; Bothmer et al. 2017). eCas9 behaved similarly to Cas9 (Figure 5A), whereas outcomes in the Cas9-TREX2 fusion protein line were markedly different from the others (Figure 5B). Cas9-TREX2 mutations were shifted towards larger deletions with frequency less than 20% (Figure 5D and E), and favoured ligation of the intact PAM-proximal side with a deletion on the PAM-distal side, in a manner not mediated by microhomology (Figure S9). This behaviour is consistent with the function of TREX2 as an exonuclease (Mazur and Perrino 2001), and the less biased set of larger deletions it generates could be beneficial in some contexts. We also tested a Cas9-2A-TREX2 construct harboring a 2A linker peptide which results in equal expression of monomeric Cas9 and TREX2; this was less effective at altering the mutational outcomes (Figure S10).

### Repair outcomes can be accurately predicted

So far, we have demonstrated that the repair outcomes are reproducible, biased, dependent on the local sequence, and mostly consistent across genetic backgrounds. These observations suggest that repair outcomes ought to be predictable from sequence alone. To test this hypothesis, we first generated candidate mutations for each gRNA within certain constraints (Methods) and derived features for each based on local sequence characteristics. We then split the set of available gRNAs into training, validation, and test sets, and trained a multi-class logistic regression model to minimize the average KL divergence between predicted and actual repair profiles (Figure 6A).

**Figure 6.**
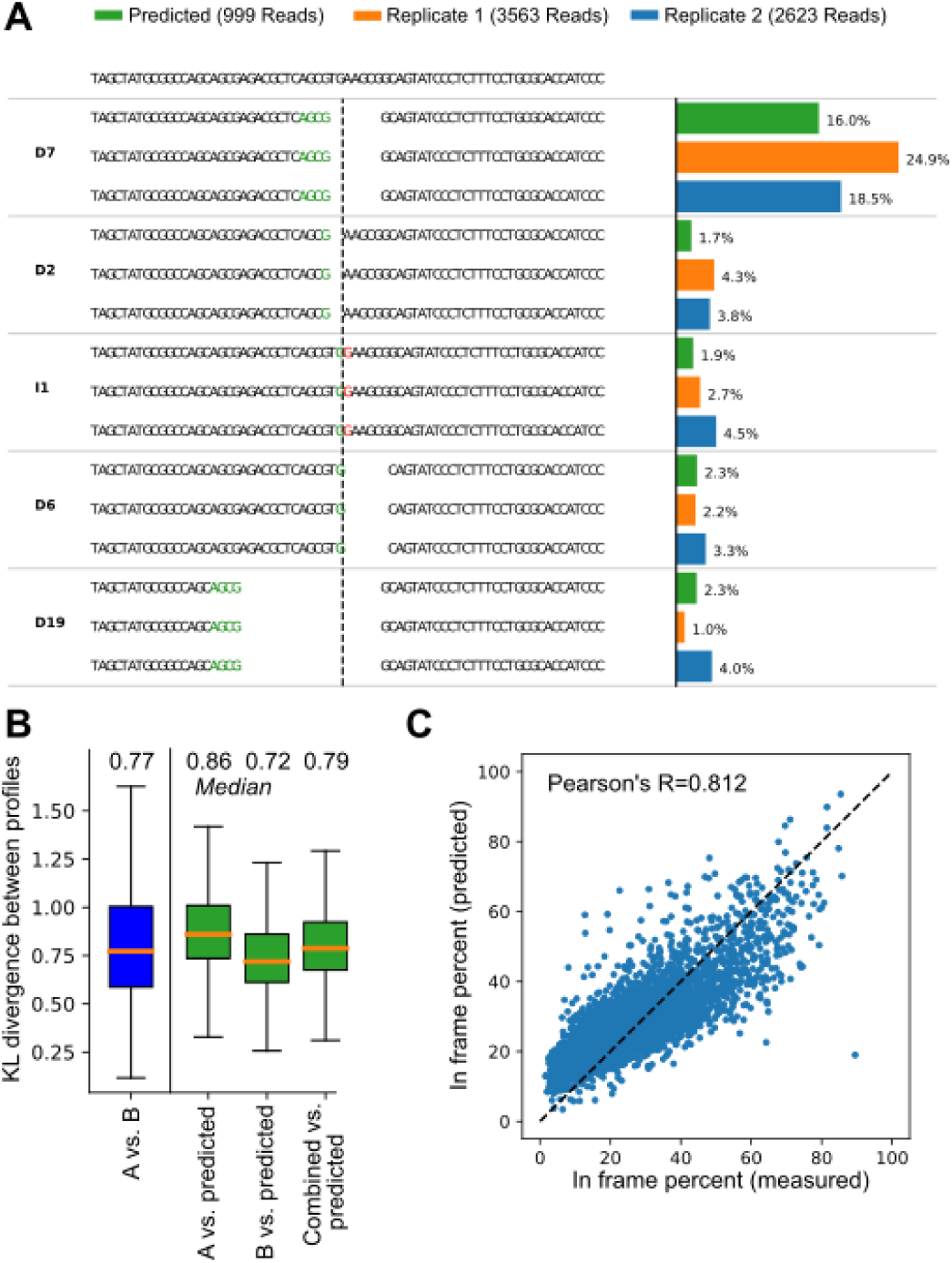
Accurate prediction of repair profiles. A.Example of a repair profile prediction with accuracy equal to the test set median (KL=0.76). DNA sequence of the target (top) is edited to produce a range of outcomes in two synthetic replicates (blue, orange bars) and one endogenous measurement (green bar). The proportions (x-axis) of the four largest mutational outcomes (e.g. “D7” - deletion of seven base pairs, “I1” - insertion of one base pair, etc.; y-axis) is consistent between the biological replicates and the prediction. Stretches of microhomology (green) and inserted sequences (red) are highlighted at the cut site (dashed vertical line). B.Repair profiles can be predicted from sequence alone. Symmetrised Kullback-Leibler divergence (KL, y-axis) between predicted and actual repair profiles (green), as well as between biological replicates A and B (blue; x-axis), with median values denoted above. Box plots: median line with median value marked, quartile box, 95% whiskers. C.Frameshift mutations can be predicted with high accuracy. Measured (x-axis) and predicted (y-axis) percent of mutations that do not produce frameshift mutations for 6,234 gRNAs.

The theoretical prediction accuracy limit is measurement repeatability. We achieved performance very close to this limit, with the average KL between predicted and measured profiles only marginally worse than that between measurements in biological replicates (0.77 vs 0.79; Figure 6B). As a consequence, we could also accurately predict the percentage of mutations that do not disrupt the reading frame (Pearson’s R=0.81 for prediction vs 0.90 for replicates) on held-out validation data (Figure 6C).

The sequence features that were most informative mirror those that we observed to induce a bias in the profiles. The features with largest influence on predictions (Table S3) contain single base insertion descriptors with positive and negative weights, recapitulating our observations that repeat of the PAM-distal T nucleotide at the cut site is particularly favoured, followed by A and then C, but that single G insertions are disfavoured, whether repeated or not. Individual deletion-related features had a weaker effect, perhaps due to their larger quantity, but substantial biases highlight the expected microhomology-related features explored above. Additional observations that warrant further study include favoring I1 insertions when there is a G in the PAM-proximal location adjacent to the cut site, and an increase in ligation of an intact PAM-distal side with a deletion on the PAM-proximal side in when a PAM-distal C is present at the cut site. While we are exploring the meaning of these biases in further work, these results demonstrate that repair outcomes can be accurately predicted from sequence alone, and in a manner that is expected to generalise to all sites in the genome. We have made this available as a webtool at https://partslab.sanger.ac.uk/FORECasT.

## Discussion

We have presented the most comprehensive study of DNA double strand break repair outcomes to date. The Cas9-generated alleles vary across sequences and between genomes, and are yet predictable near the limit of measurement repeatability. We observed three main classes of outcomes - insertions of a single base, microhomology-mediated deletions, and other deletions.

The propensities towards particular indel classes differed between cell lines and organisms, perhaps reflecting the absolute and relative activities of the corresponding repair pathways, but the particular deletions or insertions favoured by each gRNA were reproducible in K562 and consistent across lines. RPE-1 and CHO cells had elevated rates of single base insertions, while human iPSCs and mouse ESCs harboured more large insertions than other lines, and relatively more microhomology-mediated deletions. These biases can be explained by different activities of the alternative DNA repair mechanisms across lines. Stem cells (human iPSCs, mouse ESCs) have an increased rate of homology-directed repair, which shares the initial resection step with microhomology-mediated end joining (Truong et al. 2013), whereas the single based insertions favoured by other lines are likely generated by canonical end-joining. The increase in large insertions in stem cells could be explained by aberrant homology-directed repair, where strand invasion occurs in the wrong place, such that DNA synthesis before strand displacement could lead to additional sequence. While in this assay, our synthetic construct is not present on the sister chromatid to be repaired from, lower rates of mutation could be present in stem cells in general due to higher HDR activity and repeated cycles of perfect repair. However, we expect that while the mutations may happen more slowly, they would ultimately follow the patterns observed here.

As previously seen in yeast, and suggested in humans (Lemos et al. 2018) the most frequent single outcome we observed in our studies was an insertion of one base pair at the cut site, favouring repetition of the PAM-distal nucleotide in 78% of cases. This could be explained by a model where the Cas9 protein stays bound to the PAM-proximal side of the cut, and the PAM-distal end with staggered one-nucleotide overhang (Zuo and Liu 2016) is filled by DNA polymerase, and re-ligated via the non-homologous end joining pathway (Richardson et al. 2016; Lemos et al. 2018). Additionally, insertion of a thymine is greatly favoured over guanine, with a more than ten-fold increase in single base insertions if a T is present rather than a G. This could indicate either a preference of the DNA repair enzymes (especially polymerases), difference in availability of the required nucleotide triphosphate for incorporation, or propensity of Cas9 to make a staggered, rather than blunt cut when thymine is present.

Our assay was limited to confident detection of deletions of at most 30 base pairs. Recent reports indicate that substantially larger events happen at non-negligible frequency (Kosicki, Tomberg, and Bradley 2018), and indeed, some such outcomes explain the large discrepancies observed for a small number of gRNAs between our measurements and endogenous profiles. Nevertheless, in 94% of cases measured, there is good agreement between the outcomes we measured in our synthetic targets, and those at genomic targets.

Many genetic diseases, like Huntington’s or Fragile X syndrome, are due to expansions of short tandem repeats (Sutherland and Richards 1995). Such repetitive sequence serves as excellent substrate for microhomology-mediated repair, and potential correction using Cas9, especially if they also harbor a PAM site, like the CGG expansion of the *FMR2* gene in fragile XE syndrome (Gu et al. 1996). In the future, contraction of such rogue expansions could be explored as a therapy option, as the low efficacy allele replacement is not required, and simply generating a double strand break would shorten the pathogenic repeat. Indeed, a few preliminary efforts in this direction have already given promising results (Cinesi et al. 2016; Mahadevan et al. 2006; Park et al. 2015), but given the possible unintentional genomic damage (Kosicki, Tomberg, and Bradley 2018), utmost rigor is required to demonstrate safety before any applications in humans. The data and model presented here will help in guiding gRNA design towards the desired outcomes for genome-wide screens and bespoke edits.

## Methods

The code for all analyses is available at https://github.com/felicityallen/SelfTarget and also available as a runnable capsule on www.codeocean.com.

### Selection of guides and targets

We compiled our library of 41,362 total gRNA-target pairs (Table S4) from five sub-libraries aimed at testing different aspects of repair outcome generation:

- **Endogenous gRNA-Targets:** We included 86 gRNAs used in (van Overbeek et al. 2016) (“Endogenous targets”) that were compatible with our library cloning method (see below).
- **Genomic gRNA-Targets:** We selected 6,568 gRNAs from the Human v1.0 library (Tzelepis et al. 2016) as a representative sample of gRNAs used in genome-wide screening experiments. Most of these gRNAs were randomly selected from gRNAs that were also present in the GeCKOv2 library (Wang et al. 2014). Each was recorded with at least 20 mtuated reads per gRNA in each of the three K562 replicates. A subset of 3,777 gRNA-Targets, restricted to ones with over 20 reads in all samples from non-K562 cell lines, were used in comparisons across cell lines.
- **Microhomology gRNA-Targets**: We designed 28,193 guides with varying stretches, distances, and nucleotide compositions of microhomologous sequence (“Microhomology targets”). The sequences were randomly generated and then adjusted iteratively until the desired microhomology properties were achieved, ranging from targets with no microhomologies longer than 2 nucleotides within 30 nucleotides of the cut site, to targets with microhomologous sequences up to 16 nucleotides long at close range. A larger set of gRNA-targets was initially created and then filtered both for compatibility with cloning (see below), as well as to ensure each gRNA had no direct targets in the human genome (>1 mismatch between gRNA and target). Each was measured with at least 20 mtuated reads per gRNA in each of the three K562 replicates. For analysing microhomology effect in non-K562 cells, a restricted subset of 16,385 gRNA-Targets was used that also had more than 20 reads in the samples from non-K562 cell lines.
- **Microhomology Mismatch gRNA-Targets**: 294 constructs selected from Microhomology gRNA-Targets with microhomology span lengths of 6 and above were randomly altered to change one or two bases in the microhomologous sequence.
- **Conventional Scaffold gRNA-Targets**: All of the above subsets used the improved gRNA scaffold (Tzelepis et al. 2016). 6,234 gRNA-target pairs, all of which were already included in one of the first three subsets above (86 Endogenous, 3,777 Genomic (distinct set from 3,777 used for across cell line comparisons), 2371 Microhomology), were independently cloned with the alternative original gRNA scaffold. These provided independently constructed repeat measurements of the same gRNAs, with the small difference of a different scaffold (which we found to have negligible impact on mutational profile, e.g. Figure 6B).

Every target is uniquely barcoded by 10nt sequence both in the 3’ and the 5’ end (at least two mismatches between any two barcodes, randomly generated), to allow identification of each construct even in the absence of the targeted sequence. All constructs passed the filters of having no stretches of five adjacent nucleotides with at least four thymines in the gRNA sequence, since this can cause early termination of transcription; carrying BbsI restriction sites in gRNA sequence or context, including common primer sequences in gRNA sequence or context, or cutting elsewhere in the plasmid. All constructs were altered to contain a ‘G’ in the gRNA position furthest from the PAM (but not the target), for improved expression of the gRNA from the hU6 promoter.

### Construction of lentiviral library

A lentiviral gRNA expression vector lacking the scaffold pKLV2-U6(BbsI)-PKGpuro2ABFP-W was generated by removing the improved gRNA scaffold from pKLV2-U6gRNA5(BbsI)-PKGpuro2ABFP-W ((Tzelepis et al. 2016), Addgene #67974). This strategy allowed us to clone gRNA-target libraries encoding gRNAs linked to either the conventional or the improved scaffold sequences, but otherwise identical.

We generated by PCR on pKLV2-U6gRNA5(BbsI)-PKGpuro2ABFP-W two fragments encompassing the 5’ end of the AmpR cassette to U6 promoter (primers P1-P2, Table S1) and PGK promoter to the 3’ end of the AmpR cassette (primers P3-P4), respectively. Primer overhangs were designed to generate overlapping ends and pKLV2-U6(BbsI)-PKGpuro2ABFP-W was obtained by Gibson assembly (NEBuilder HiFi DNA Assembly Master Mix, NEB) of the two fragments. BbsI restriction sites were present downstream from the U6 promoter for subsequent cloning of the gRNA-target library inserts.

Library cloning started by PCR amplification of the 170 nt oligonucleotide pool of designed sequences (CustomArray) encoding gRNA and target sequence, separated by a spacer harbouring two BbsI restriction sites (Supplementary Figure 1). Primer pairs P5-P6 and P7-P8 (Table S1) were used to amplify oligos compatible with the conventional or improved scaffold respectively. Gibson assembly (Gibson 2011) was employed to fuse the amplified pool to a 193 nt G-block fragment (IDT) encoding either a conventional or improved version of the gRNA scaffold (Tzelepis et al. 2016) and a spacer. 1:1 molar ratio was mixed in three reactions incubated 1 h at 50°C and subsequently pooled. The resulting 318 bp circular DNA was column purified (PCR purification kit, QIAGEN) and treated with Plasmid-Safe ATP-Dependent DNase (Epicentre) to remove linear DNA, followed by linearisation with BbsI at 37°C for 2 h. The resulting 296 bp linear fragment was ligated into scaffold-less pKLV2-U6(BbsI)-PKGpuro2ABFP-W. Ligations (T4 DNA Ligase, NEB) were performed in triplicate, pooled and used in up to 10 electroporation reactions to maximise library complexity. Next generation sequencing verified that libraries encoded 12,113 and 48,412 gRNA-target pairs for the conventional and improved scaffold, respectively.

### Generating the TREX2 construct

The Cas9-TREX2 and Cas9-2A-TREX2 vectors were made by fusing the human TREX2 open reading frame (GBlock, IDT) to the C-terminus of the Cas9 protein (Cong et al. 2013) with a GGGS linker or an intervening T2A peptide. These were cloned by Gibson assembly into a piggyBac vector (pKLV-Cas9), driven by a EFS promoter, and containing a blasticidin selectable marker. Genbank files of the final vectors are provided in the Supplementary Information.

### Cell culture

K562, K562-Cas9 (kind gift by Etienne De Braekeleer) and all K562-derived lines (see below) were cultured in RPMI supplemented with 10% FCS, 2 mM L-glutamine, 100 U/ml penicillin and 100 mg/ml streptomycin. CHO-Cas9 and 293FT (Invitrogen) cells were cultured in Advanced DMEM supplemented with 10% FCS, 2 mM L-glutamine, 100 U/ml penicillin and 100 mg/ml streptomycin. HAP1-Cas9 were cultured in IMDM supplemented with 10% FCS, 2 mM L-glutamine, 100 U/ml penicillin and 100 mg/ml streptomycin. RPE-1-Cas9 cells were cultured in DMEM:F12 supplemented with 10% FCS, 0.26% sodium bicarbonate, 2 mM L-glutamine, 100 U/ml penicillin and 100 mg/ml streptomycin.

E14TG2a mouse ES cells supplied by Dr Meng Li (Cambridge Stem Cell Institute) were cultured in High glucose DMEM supplemented with 15% FBS, 2 mM L-glutamine, 0.1 mM 2-mercaptoethanol and 1,000 U/ml leukemia inhibitory factor (LIF; Millipore).

iPSCs (REC 15/WM/0276) were cultured in vitronectin (Life Technologies Ltd.) coated plates and TeSR-E8 medium (Stemcell Technologies). E8 media was changed daily throughout expansion and all experiments. All cell lines were cultured at 37°C, 5% CO_2_.

### Lentivirus production and transduction of cell lines

Supernatants containing lentiviral particles were produced by transient transfection of 293FT cells using Lipofectamine LTX (Invitrogen). 5.4 µg of a lentiviral vector, 5.4 µg of psPax2 (Addgene #12260), 1.2 µg of pMD2.G (Addgene #12259) and 12 µl of PLUS reagent were added to 3 ml of OPTI-MEM and incubated for 5 min at room temperature. 36 µl of the LTX reagent was then added to this mixture and further incubated for 30 min at room temperature. The transfection complex was added to 80% confluent 293FT cells in a 10-cm dish containing 10 ml of culture medium. After 48 h viral supernatant was harvested and stored at –80 °C. Fresh medium was added and lentiviral supernatant was collected a second time 24 h later. When necessary we prepared larger amounts of lentivirus by scaling up the procedure above.

For lentiviral transduction K562 and K562-derived cells (see below), CHO-Cas9, HAP1-Cas9 and RPE-1-Cas9 cell lines were incubated with the lentiviral supernatant in a single cell suspension in presence of 8 µg/ml polybrene (Hexadimethrine bromide, Sigma) followed by centrifugation for 30 min at 1,000xg. E14TG2a mouse ESCs transduction was performed incubating cells in suspension for 30 min in presence of 8µg/ml polybrene. iPSC transduction was performed in a single cell suspension obtained by incubating cells with Accutase for 10 min (Millipore Corporation) and cells were plated in E8 medium supplemented with 8µg/ml polybrene.

### Generation of K562-eCas9, K562-Cas9-TREX2, K562-Cas9-2A-TREX2 and E14TG2a-Cas9 lines

Cell lines stably expressing Cas9 were generated by lentiviral transduction followed by selection in the presence of Blasticidin (Cambridge Bioscience) to ensure high Cas9 activity. K562 cells were transduced using eSpCas9(1.1) (Slaymaker et al. 2016) (Addgene #71814), Cas9-TREX2 and Cas9-2A-TREX2 vectors to generate K562-eCas9, K562-Cas9-TREX2 and K562-Cas9-2A-TREX2 respectively. E14TG2a-Cas9 cells were generated by transducing E14TG2a cells with pKLV2-EF1aBsd2ACas9-W (Tzelepis et al. 2016)(Addgene #67978).

### Screening and sequencing of repair outcomes

Cell lines were infected at a multiplicity of infection (MOI) ranging from 0.5 to 0.6 and at a coverage ranging from 500X to 1600X. Total number of cells, MOI and coverage for each screen are listed in Table S2. For each line, at least two separate infections were performed and treated separately as biological replicates. 24 h after transduction (72 h for iPSCs) puromycin was applied to the culture medium to select for successfully transduced cells and maintained throughout the screen. Cells were cultured for a minimum of 7 days and up to 20 days in the case of K562 and K562-Cas9 cells. Enough cells were passaged and collected to maintain coverage higher than at the time of infection.

Upon collection cells were centrifuged and pellets were stored at −20°C prior extraction of genomic DNA. Briefly, cell pellets were resuspended into 100 mM Tris-HCl pH 8.0, 5 mM EDTA, 200 mM Nacl, 0.2% SDS and 1 mg/ml Proteinase K and incubated at 55°C for 16 h. DNA was extracted by adding 1 volume of isopropanol followed by spooling, washed twice in 70% ethanol, centrifuged and resuspended into TE.

For sequencing, the region containing the target surrounded by the context was amplified by PCR using primers P10-P12 or P11-P12 respectively for the conventional and improved scaffold (Table S1) with Q5 Hot Start High-Fidelity 2× Master Mix (NEB) with the following conditions: 98 °C for 30 s, 24 cycles of 98 °C for 10 s, 61 °C for 15 s and 72 °C for 20 s, and the final extension, 72 °C for 2 min. Alternatively, both gRNA and target were amplified using primers P9-P12. For each gDNA sample the amount of input template was calculated keeping into account coverage and the amount of gDNA per single cell depending on the species and the ploidy of each line, and PCR reactions were scaled up accordingly. The PCR products were pooled in each group and purified using QIAquick PCR Purification Kit (QIAGEN). Sequencing adaptors were added by PCR enrichment of 1 ng of the purified PCR products using forward primer P13 and indexing reverse primer P14 with KAPA HiFi HotStart ReadyMix with the following conditions: 98 °C for 30 s, 12-16 cycles of 98 °C for 10 s, 66 °C for 15 s and 72 °C for 20 s, and the final extension, 72 °C for 5 min. The PCR products were purified with Agencourt AMPure XP beads, quantified and sequenced on Illumina HiSeq2500 or HiSeq4000 by 75-bp paired-end sequencing using the following custom primers: P15-P18 for sequencing of both gRNA and target, P16-P18 (conventional scaffold) or P17-P18 (improved scaffold) for target-only sequencing.

### Sequence analysis

We processed the generated sequence data to compile repair profiles as follows. First, we combined the partially overlapping paired-end reads into a single sequence using pear v.0.9.10 (Zhang et al. 2014) with options “-n 20 -p 0.1” (minimum combined sequence length of 20, probability of no overlap below 0.1). To assign reads to constructs, we required that at least one of the unique 3’ and 5’ barcodes to be present with at most one mutation, and confirm that the read can be aligned to the template in such a way that at least 80% of the read characters has a match in the construct template (i.e. there can be a large deletion in the read around the cut site, but minimal misalignment outside that region). The alignment was done using a custom dynamic programming algorithm in which the two sides of the cut site are independently aligned and then efficiently combined. This algorithm allows large deletions at a specified place (the expected Cas9 cut site) without penalty, while imposing substantial gap penalties elsewhere, and unlike generic tools, works for relatively short sequences.

We first used this approach on sequences of the plasmid library to compile a set of null mutations that are present before any editing experiments. These null mutations are then used as templates for alignment of sequencing reads from the screens, again using the custom dynamic program, so that mutations already existing in the plasmid library (due to oligo synthesis errors or somatic mutation) are not erroneously attributed to Cas9 activity.

### Repair profile analysis

We store the repair profile as a collection of read counts per indel, where each indel is characterised by its size, type and location with respect to the cut site. To calculate similarity of repair profiles, we use the symmetrized Kullback-Leibler divergence (“KL”). Standard Kullback-Leibler divergence is calculated as

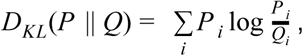

where *i* indexes the different indels, and *P* _*i*_ and *Q*_*i*_ are their normalized proportion of total mutated read counts in the compared samples; we employ the symmetric form,

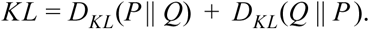

To avoid division by zero, we add small pseudocounts of 0.5 to all indels present in one sample, but not the other, in the computation.

We classified all indels into types (insertion or deletion), size (at least 4 nt or less than 4 nt), and likely causal mechanism of the event. We classified deletions as likely generated by microhomology if at least 2nt of matching sequence were present on either side of the cut, and likely not generated by microhomology otherwise. To analyse indel prevalence by size, we classified indels into deletions of 1-30 nt and insertions of 1 to 10 nt. In this case, information about deletions larger than 30 and insertions larger that 10 nt was excluded from the analysis since they are not well-detected by our method. In this, and other results presenting accumulated measurements across gRNAs (Figure 3A,3B,3C,5A) we first normalized all indel counts for each gRNA by the total number of mutated reads for that gRNA, such that all gRNAs weigh equally towards each measurement, rather than proportionally to their read coverage.

### Predicting repair profiles

For each gRNA-target pair we generated candidate indels by considering all possible insertions up to size 2, within 3 nucleotides of the cut site, and all deletions spanning the cut site with a left edge from one right of the cut site to up to 30 nucleotides left of the cut site and a right edge up to 30 nucleotides the other way, up to a maximum deletion size of 30. For each of these candidates, we computed a set of 2,137 binary features that describe the length, location and nucleotide composition of inserted sequences, microhomologies and their neighbouring nucleotides. Many of these features also comprise pairwise ‘AND’ results between features to capture interaction effects. We modelled the probability of each possible outcome using a logistic that ensures the sum of all possible outcomes for a give gRNA sums to one. i.e. the probability of the jth mutation for a given gRNA is modelled as

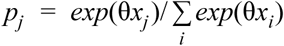

where θ is the vector of parameter weights, and *x*_*j*_ is the feature vector for that mutation; the sum is over all mutations for a particular gRNA. We then minimize the L2-regularized, non-symmetric KL divergence of these probabilities when compared to the measured proportions, by computing closed form partial gradients with respect to the kth parameter θ_*k*_ and using L-BFGS-B within scipy.optimize.minimize to perform gradient descent optimization of this metric (Zhu et al. 1997; Jones, Oliphant, and Peterson 2001).

For development of the predictor, we randomly selected gRNA-target pairs from the “Microhomology gRNA Targets” set, restricted to those with more than 100 reads in K562 cells, and without a corresponding counterpart in the “Conventional scaffold gRNA Targets” set, and performed 2-fold cross-validation training and hyperparameter tuning. With just 1000 training examples, the training and test scores converged for this feature set with a regularization constant of 0.01, so the parameters trained with these settings were selected for further validation (Figure S11). We used these trained parameters to predict profiles by applying the associated probabilities to generate 1000 counts each for all gRNA-target pairs in the “Conventional scaffold gRNA Targets” set (dropping indels predicted to have less than 1 count). We then validated the accuracy of these profiles by comparing against measurements for these gRNA-target pairs in both the Conventional Scaffold library and for its corresponding counterparts in the other subsets with Improved Scaffold.

## Author Contributions

FA: designed experiments, analysed data, wrote paper. LC: designed experiments, performed experiments, wrote paper. CA: performed experiments in human iPSCs. AJS, EM: performed experiments in mouse ESCs. VK: analysed data, wrote paper. MO: analysed data. PDA, PP: performed experiments. MK, ARB: generated TREX2 constructs. HH: generated CHO-Cas9 line. YG, FM-M, SPJ: generated RPE-1-Cas9 and HAP1-Cas9 lines. LP: designed experiments, wrote paper. All authors contributed to drafting the manuscript.

## Acknowledgements

FA was supported by a Royal Commission for the exhibition of 1851 Research Fellowship. LP was supported by Wellcome, and Estonian Research Council (IUT 34-4). HPH was supported Supported by a Wellcome Trust grant (Wellcome 200848/Z/16/Z) and a Wellcome Trust Strategic Award to the Cambridge Institute for Medical Research (Wellcome 100140). YG is funded by Cancer Research UK, C6/A18796 and a Wellcome Trust Investigator Award 206388/Z/17/Z in the Jackson lab. FMM was funded by a Marie Curie Intra-European Fellowship, project number 626375, DDR SYNVIA, and by a Wellcome Trust Investigator Award 206388/Z/17/Z and AstraZeneca Collaborative Award in the Jackson lab. We thank Jana Eliasova for help with Figure 1, and Andrew Lawson for comments on the text.

